# SOGO-SOFI, light-modulated super-resolution optical fluctuation imaging using only 20 raw frames for high-fidelity reconstruction

**DOI:** 10.1101/2022.09.03.506455

**Authors:** Fudong Xue, Wenting He, Dingming Peng, Hui You, Mingshu Zhang, Pingyong Xu

## Abstract

Taking advantage of the stochastic photoswitching of genetically encodable reversibly photoswitchable fluorescent proteins (RSFPs), super-resolution optical fluctuation imaging (SOFI) and its variant photochromic stochastic optical fluctuation imaging (pcSOFI) are valuable tools for wide field super-resolution (SR) imaging. Live-cell (pc)SOFI, which requires a small number of original frames to reconstruct an SR image, is prone to structural discontinuity artifacts and low spatial resolution. Herein, we developed a repeated synchronized on- and gradually off-switching SOFI (SOGO-SOFI) that maximized the photoswitching frequency of RSFPs by light modulation and required only 20 frames for high-quality reconstruction. Live-cell SOGO-SOFI imaging of the endoplasmic reticulum (ER) exhibited 10 times higher temporal resolution (100 fps) and fewer artifacts than pcSOFI. Moreover, a combination of SOGO-SOFI with Airyscan further increased the image contrast and the resolution of Airyscan by a factor of 1.5 from 140 nm to 91 nm. The capabilities of SOGO-SOFI were further demonstrated by dual-color imaging of nucleolar proteins in mammalian cells and deep imaging of ER structures in thick brain slices (20.6 µm).

## Introduction

Photo-switchable fluorescent proteins (RSFPs) are fluorescent proteins that can be switched on and off by lights at different wavelengths in a reversible manner. Using the different fluorescence states (on or off) of RSFPs at different sampling time points, the temporal or spatial correlation analysis of the fluorescent signal of each pixel can significantly improve the signal-to-noise ratio and contrast of the image. For example, using an RSFP Dronpa, optical lock-in detection (OLID) can separate signaling molecules from the high-fluorescence background by correlating time-lapse images of different fluorescence intensities over time[1]. However, OLID imaging uses optical modulation to allow all fluorescent molecules in the diffraction range to fluoresce at the same time or to be turned off at the same time, resulting in insignificant fluctuations in the fluorescence signal at successive sampling time points on a single pixel. Therefore, OLID cannot break the optical diffraction limit and improve spatial resolution. The super-resolution optical fluctuation imaging (SOFI) technique also performs fluorescence fluctuation correlation analysis on time-lapse images of RSFPs. However, unlike OLID, it allows different RSFPs or blinking molecules within the diffraction limit to randomly appear and fluctuate by adjusting the energy of the excitation light. Therefore, SOFI can improve spatial resolution by calculating the higher-order cumulant of temporal fluorescence fluctuations recorded in a sequence of images[2-4].

SOFI generally offers a 2–4-fold spatial resolution enhancement beyond the diffraction limit. It can also enhance signal contrast by removing the background fluorescence, which shows much less fluctuation than that of the signal when analyzing the temporal and spatial correlations of fluorescence on pixels. Furthermore, SOFI is a very flexible method that can be easily integrated into other imaging modalities. For example, spinning disk confocal microscopy was employed in association with SOFI to obtain fast 3D super-resolution (SR) images[5]. Taking advantage of the complementarity between photo-activation localization microscopy (PALM)[6] and SOFI, unprecedented functional exploration of focal adhesions was demonstrated[7]. Combing the two-photon light-sheet illumination with SOFI provided a promising way to realize long-term, deep-tissue SR 3D imaging[8]. The hybridization of SOFI with structured illumination or image scanning microscopy, including Airyscan, was reported to increase spatial resolution[9-11].

SOFI imposes no constraint on the type of probe used, as long as the probe exhibits robust fluorescence fluctuation. The fluctuation could be attributed to fluorescence intermittency intrinsic to the probe, such as quantum dots[2], or redox buffer assisted photoblinking of the probe, such as organic fluorophores[3]. However, because of either the specificity of labeling or the reducing nature of the buffer, these probes are not suitable for imaging living systems. Taking advantage of genetically encodable RSFPs[12-14], photochromic stochastic optical fluctuation imaging (pcSOFI)[12] was evolved from SOFI to enable live-cell compatible SR imaging. Under certain illumination conditions, the ratio of the on-switching molecules to the off-switching molecules of RSFPs in a labeled sample will eventually reach equilibrium. As a result, the overall brightness of the sample stays constant, but stochastic fluorescence intensity fluctuations in each pixel of the camera can be observed. In conventional pcSOFI, time-lapse images are acquired when the RSFP ensemble is at equilibrium, and the independent stochastic photoswitching of RSFPs is utilized for analysis.

Compared to PALM and stochastic optical reconstruction microscopy (STORM)[15], pcSOFI has increased temporal resolution and is more favorable for live imaging, albeit with inferior spatial resolution. Compared to structured illumination microscopy (SIM)[16], nonlinear structured illumination microscopy (NL-SIM)[13, 17, 18], stimulated emission depletion (STED) microscopy[19], and reversible saturable optical fluorescence transitions (RESOLFT)[20], pcSOFI has comparable spatial resolution and requires no complicated optical equipment or expertise. However, pcSOFI uses the cumulant of fluorescence fluctuations per pixel for SR reconstruction. If insufficient fluorescence fluctuations are recorded at some structured positions, the final reconstructed SR image will lose structures at these positions and display discontinuity artifacts. Another disadvantage of pcSOFI is that its temporal resolution is still impractical for fast real-time imaging, as 500–2000 frames are often used to reconstruct an SR image with acceptable fidelity[2, 7, 21, 22], which limits the application of SOFI and SOFI-combined SR imaging in living cells. Reducing the number of raw frames for reconstruction can improve the temporal resolution of pcSOFI, but at the expense of the accumulation of fluctuation, resulting in reduced spatial resolution, loss of information, and discontinuous artifacts.

Several methods have been developed to increase the temporal resolution of SOFI. Using a strategy to label the same target by three types of quantum dots (QDs) with distinct emission spectra, joint-tagging SOFI (JT-SOFI) collects 100 frames per channel simultaneously for SOFI reconstruction, enabling an increased labeling density as well as a temporal resolution of 4.5 s[23]. However, this method is limited by few choices of spectrally separated fluorescent probes and is not suitable for two-color imaging. A modified SOFI algorithm capable of eliminating the fluorescent background and readout noise was reported to reduce the number of raw frames to 25 and achieve a temporal resolution of 1.5 s[24]. Nevertheless, clear structural discontinuity was found in the microtubule resolved by this algorithm, raising doubts about the reliability of the method. Through engineering QDs with enhanced blinking, real-time SOFI variance imaging with only 10 frames was demonstrated[25]. However, the broad application of this method is hampered by the requirement for tailor-made probes and limited target diversity of QDs in living system. Through carefully optimizing the frame rate and the illumination conditions to match the properties of an RSFP Dreiklang, a 1.5 s temporal resolution was realized by pcSOFI[26]. However, at least 500 raw images were required to obtain the final image with an acceptable SNR. Balanced SOFI (bSOFI) can solve the structural discontinuity problem to a certain extent[4]. However, b-SOFI is a post-processing algorithm based on the result of SOFI reconstruction, and the lost weak signal still cannot be retrieved.

Therefore, there is a need to balance the improvement of temporal resolution and the problem of structural discontinuity when performing SOFI. We propose a new method to generate changes in fluorescence images by modulating the optical switch of RSFP, thereby generating fluorescence fluctuations, to improve the fluorescence fluctuation signals on a unit pixel per unit time and increase the resolution of pcSOFI. Our strategy is to first modulate all fluorescent molecules to the on state and then turn them off asynchronously in sequence during the recording of the fluorescent signal, resulting in a time series of images with different numbers and localization distributions of fluorescent molecules. Time-dependent changes in the distribution of fluorescent molecules produce large fluorescence fluctuations, which can significantly increase the accumulation of fluctuation on a single pixel per unit time, thereby improving the temporal resolution of SOFI and reducing discontinuities in the final reconstructed SR image. We denoted this method as repeated synchronized <u>o</u>n- and <u>g</u>radually asynchronous <u>o</u>ff-switching <u>SOFI</u> (SOGO-SOFI).

In this study, we first verified the feasibility of SOGO-SOFI using both simulated data and biological experiments. Then, fast live imaging was performed to prove the utility of the method. Furthermore, as a demonstration of the flexibility of the method, we combined SOGO-SOFI with Airyscan microscopy and exemplified the application of this fusion method for deep SR imaging of cell nuclei and brain slices.

## Result Simulations

Two sets of simulated data of SOFI and SOGO-SOFI were produced by the SOFI Simulation Tool[27]. The photoswtiching contrast and the off-switching kinetics of a monomeric green RSFP, Skylan-S[13], were experimentally determined and input as parameters for data simulation (Supplementary Note). For SOFI, the data has a constant number of single molecules per frame and *N* frames in total. For SOGO- SOFI, while the total number of frames was the same, the number of single molecules per frame was cycled between a fixed value (depending on the ratio of fluorescent molecules in the ON state to all fluorescent molecules, and on the illumination power of 405 nm laser) and a minimum value (corresponding to the OFF state) every *F* frames, and within each cycle, the number was exponentially decreased according to the off-switching kinetics of Skylan-S (Figure 1a–c). We first test the effect of different raw frame numbers on the quality of the final reconstructed SR image. Reducing the number of raw frames significantly reduced the quality of the reconstructed SOFI image and created artifacts of missing structures and discontinuities (Supplementary Figure 1). Increasing the ratio of ON molecules to all fluorescent molecules by SOGO-SOFI modulation can significantly reduce the structural discontinuity generated during reconstruction with a small number of frames (Supplementary Figure 1). In order not to produce structural discontinuities visible to the naked eye, at least 20 original images are required for reconstruction with SOGO-SOFI, and all fluorescent molecules need to be modulated to the ON state (Supplementary Figure 1). Therefore, to meet the requirement of high temporal resolution in live-cell imaging as much as possible, in the subsequent simulations and biological experiments, we choose the maximum activated fluorescent molecules and 20 raw frames for SOGO-SOFI. The fluorescence intensity fluctuation of the SOGO-SOFI data was much higher than that of the SOFI data in a representative pixel (Figure 1d), indicating a larger SOFI signal and a higher signal-to-noise ratio of SOGO-SOFI. In SOGO-SOFI, the number of blinking molecules exhibited periodical rise and fall along the imaging time course, whereas in SOFI, it stayed constant at a remarkably lower level than that in SOGO-SOFI (Figure 1e), suggesting that SOGO-SOFI provided higher labeling density than SOFI.

**Figure 1.**
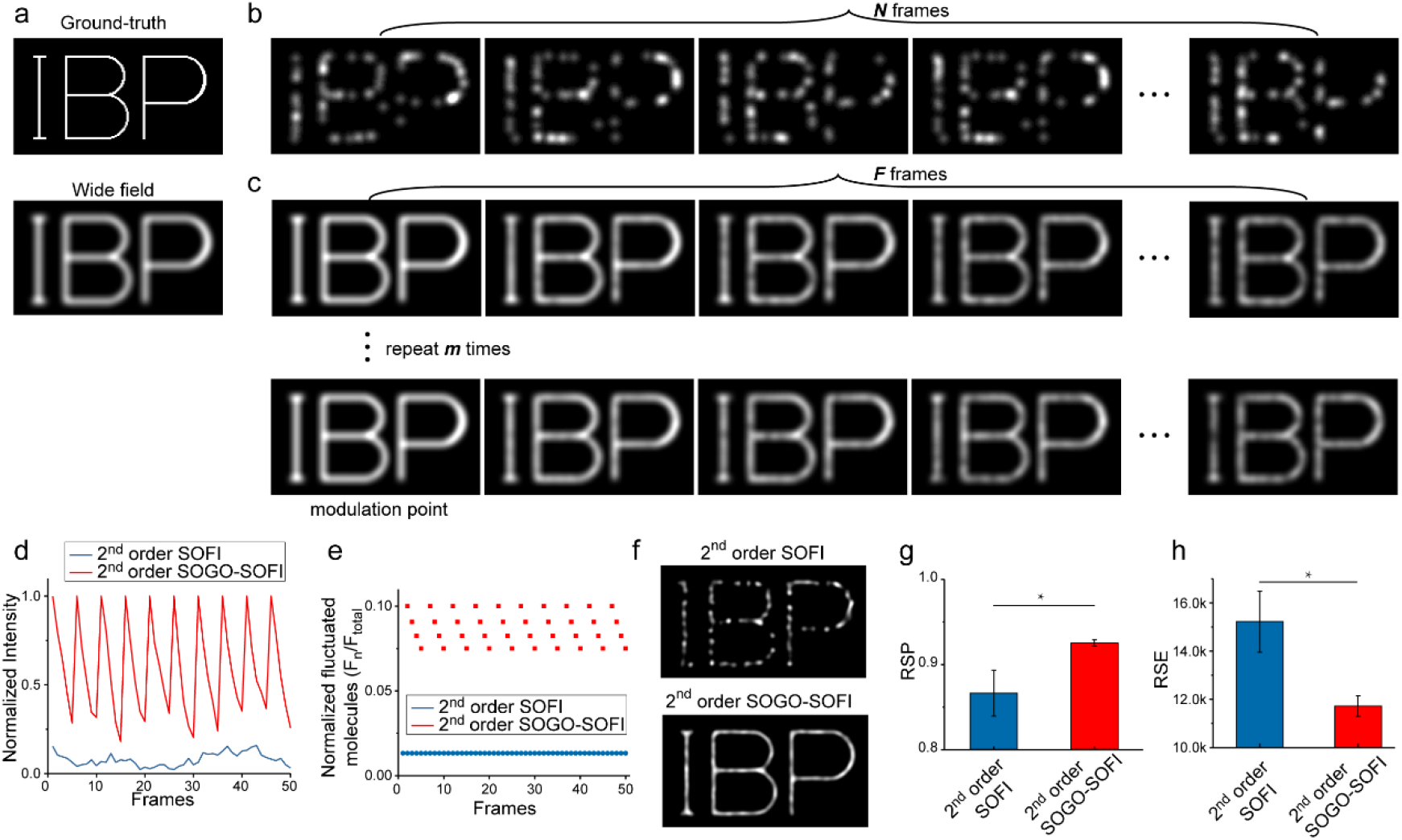
Comparison of SOGO-SOFI and SOFI by simulation. (a) Top: Ground-truth image; Bottom: Sum of 20 raw frames as the wide field image. (b) Consecutive raw images for SOFI. (c) Consecutive raw images for SOGO-SOFI. (d) Normalized fluorescent intensity trace of one representative pixel in (b) and (c). (e) Ratio of blinking molecules to total molecules in SOFI and SOGO-SOFI raw data. (f) Top: 2^nd^ order SOFI image using 20 raw frames; Bottom: 2^nd^ order SOGO-SOFI image using 20 raw frames. (g) RSP between the wide field image and the SR images. (h) RSE between the wide field image and the SR images. Data are reported as mean ± SD. All p values were calculated by the two-tailed Student’s *t*- test, *n* = 3, * indicates p < 0.05.

To quantitatively assess the quality and artifacts of reconstructed SR images from different raw frames, we used the SQUIRREL analysis[28]. To be specific, the resolution-scaled Pearson coefficient (RSP) and the resolution-scaled error (RSE) were calculated to present the Pearson correlation coefficient and the root-mean-square error between the reference (the wide field image in Figure 1a) and the resolution-scaled image. The higher the RSP value and the lower the RSE value, the higher the image quality of the reconstructed SR image. An error map was also generated by calculating the pixel-wise absolute difference between the reference and resolution-scaled image. We found that the frame number of 20 was sufficient for SOGO-SOFI reconstruction with an acceptable quality as compared to a frame number of 2000 (Supplementary Figure 2). In contrast, SOFI requires at least 200 raw frames to achieve the same effect of reconstructed images with 20 raw frames by SOGO-SOFI (Supplementary Figure 2b). Intuitively, the SR image reconstructed from 20 raw frames by SOGO-SOFI showed much higher structure continuity than that reconstructed by SOFI with the same raw frames (Figure 1f). The RSP of the SOGO-SOFI image was notably higher and the RSE was notably lower than those of the SOFI image (Figure 1g & h).

### Validation of SOGO-SOFI in mammalian cells

The simulation results show clearly that SOGO-SOFI can effectively reduce the structural loss and discontinuity artifacts caused by the small number of frames during SOGO reconstruction. To demonstrate the SOGO-SOFI’s effect experimentally, we transfected U-2 OS cells with Lifeact-Skylan- S fusions. Images were acquired under total internal reflection fluorescence (TIRF) microscopy using the imaging schemes illustrated in Supplementary Figure 3. Briefly, for SOFI, only a continuous 488- nm laser was used; for SOGO-SOFI, an additional 405-nm laser pulse was applied every 25 ms for 5 ms. The datasets acquired by SOFI and SOGO-SOFI each contained 50 frames in total and were reconstructed by the bSOFI algorithm[4]. Remarkably, the SR images of SOGO-SOFI showed much better structural continuity than those of SOFI (Figure 2a–c). As shown in Figure 2c (arrows), SOGO- SOFI has a much stronger ability to extract weak signals that cannot be resolved by SOFI (Figure 2b, arrows). Furthermore, two adjacent actin filaments could be observed by SOGO-SOFI but not by SOFI (Figure 2d, arrow heads), indicating that SOGO-SOFI had higher spatial resolution than SOFI using the same 50 frames for reconstruction. The spatial resolution of SOGO-SOFI, as indicated by the distance between the two distinguishable filaments, was 163 nm (Figure 2e). We also compared the RSP and the RSE metrics of SOGO-SOFI and SOFI. As expected, the SR image of SOGO-SOFI had prominently higher quality and fewer artifacts than that of SOFI (Figure 2f–g).

**Figure 2.**
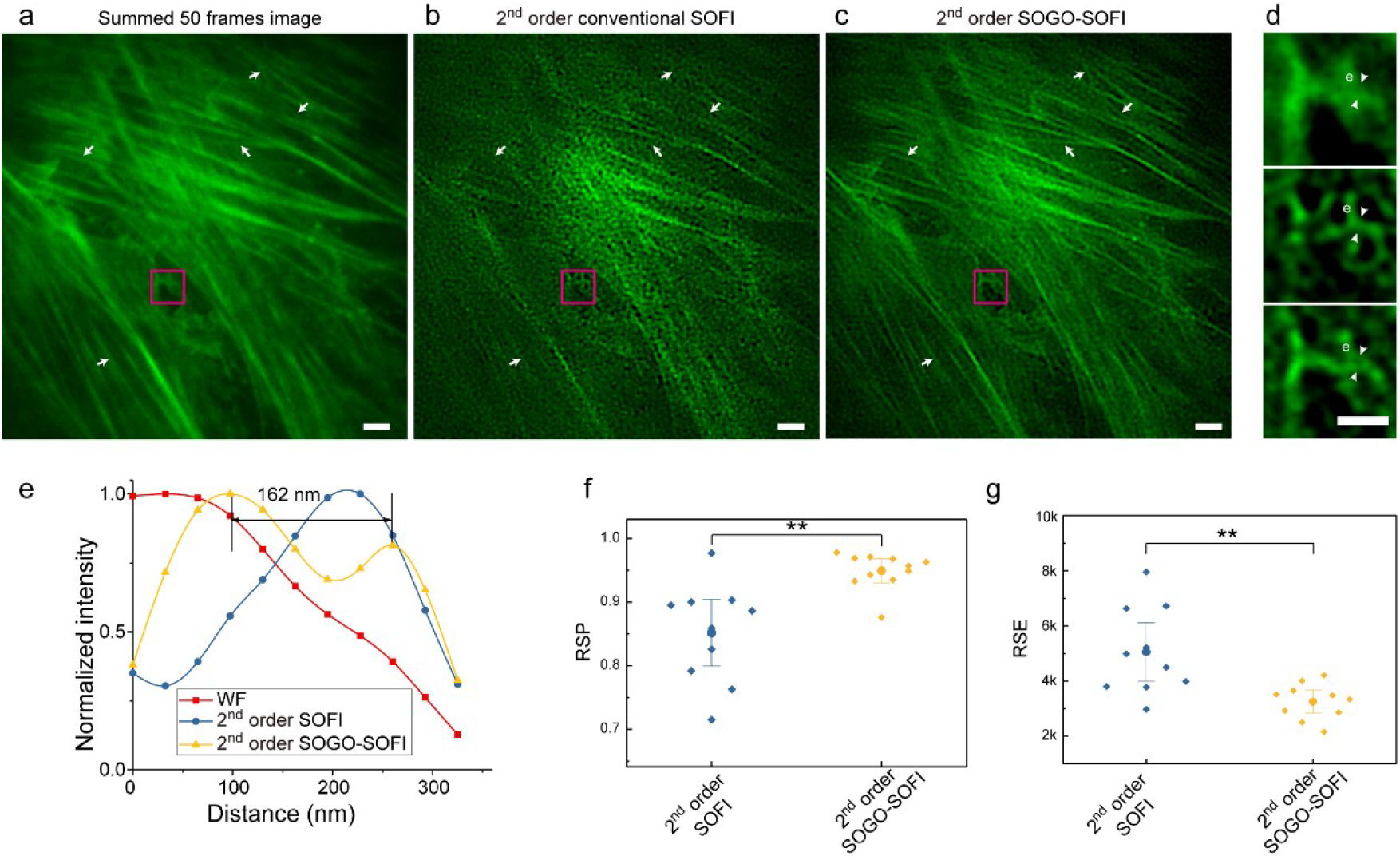
Validation and quality assessment of SOGO-SOFI by imaging F-actin of U-2 OS cells. (a) Sum of raw TIRF images as the wide field (WF) image. (b) Second-order SOFI SR image. (c) Second-order SOGO-SOFI SR image. (d) Zoomed-in images inside the magenta boxes in (a–c). (e) Intensity profiles along the lines indicated by the arrows in (d). (f) RSP between the second-order conventional SOFI and SOGO-SOFI SR images. (g) RSE between the second-order conventional SOFI and SOGO- SOFI SR images. Scale bars: 2 μm in (a–c) and 1 μm in (d). Data are reported as mean ± SD. All p values were calculated by the two-tailed Student’s *t*-test, *n* = 10, ** indicates p<0.001.

### SOGO-SOFI reveals fast SR dynamics of mitochondria and endoplasmic reticulum network

To demonstrate the utility of SOGO-SOFI for fast live imaging, we transfected COS-7 cells with Skylan- S-labeled TOM20, an outer mitochondrial membrane receptor. Compared to the wide field image, SOGO-SOFI more clearly resolved the outer membrane structure of the mitochondria (Figure 3a). Moreover, the dynamic movement of the TOM20 was captured by SOGO-SOFI at 10 Hz (Figure 3b). The endoplasmic reticulum (ER) is the largest intracellular organelle and forms a complex network of dense matrices and well-separated tubules. Insufficient spatiotemporal resolution will lead to misinterpretation of the structure of the ER dense matrices[29, 30]. Therefore, to verify the excellent performance of SOGO-SOFI, we imaged COS-7 cells expressing calnexin-Skylan-S fusions. Remarkably, SOGO-SOFI resolved the ER dense tubular structure that appeared as a continuous sheet in the WF image (Figure 3c). Notably, using only 20 raw frames (100 Hz), SOGO-SOFI successfully revealed the rearrangement of the inside fistulous structures in the dense tubular matrices (Figure 3d). In addition, SOGO-SOFI also allowed us to capture the continuous and rapid changes of the large-view ER tubular network (Figure 3e).

**Figure 3.**
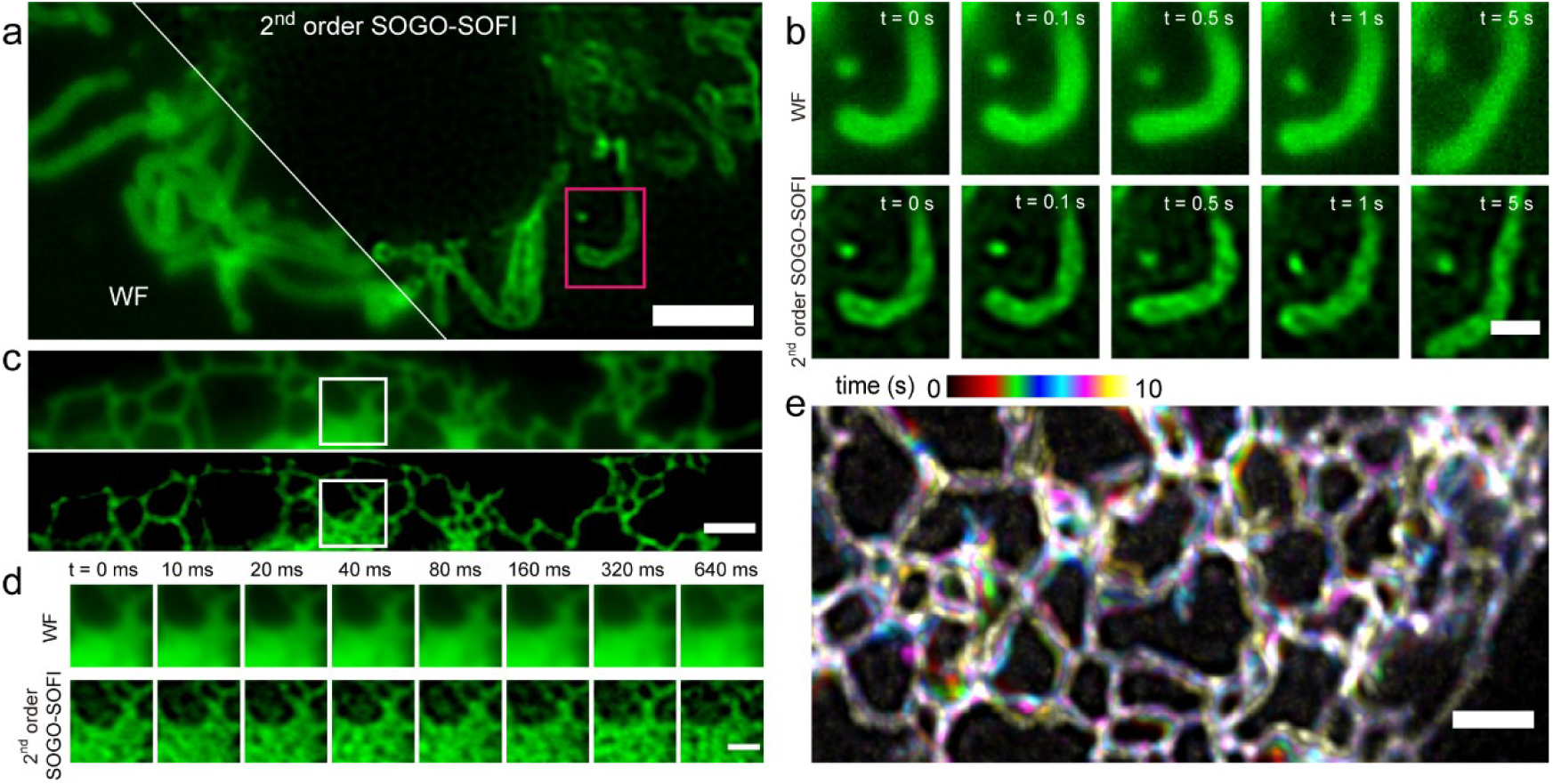
Fast live imaging of mitochondria and ER by SOGO-SOFI. (a) Wide field (WF) and 2^nd^ order SOGO-SOFI SR image of Skylan-S labeled mitochondrial outer membrane protein TOM20 in live COS-7 cell. (b) Magnified view of the time-lapse images of the boxed region in (a). (c) WF and 2^nd^ order SOGO-SOFI SR image of Skylan-S labeled ER protein calnexin in live COS-7 cell. (d) Magnified view of the time-lapse images of the boxed region in (c). (e) Motion map of the large-view ER tubular network recorded at 40 Hz over 10 s, color coded by time. Scale bars: 2 μm in (a, c, e), 1 μm in (b, d).

### Combining SOGO-SOFI with Airyscan

Like SOFI, SOGO-SOFI could be easily merged with other imaging techniques to gain combined advantages. Airyscan exploits the pixel-reassignment principle to increase the spatial resolution and the signal-to-noise ratio by 1.7× and 4-8× respectively[31]. Technically, Airyscan is a combination of the confocal microscopy, the Airyscan detector and a linear deconvolution, and thus is ideal for deep imaging. We proposed that a fusion method of SOGO-SOFI and Airyscan (we termed SS-Airyscan) would offer additive enhancements of spatial resolution and image contrast and thus provide advantages for deep SR imaging.

We adopted a fusion strategy for SS-Airyscan that was different from previous reports combining SOFI with Airyscan[9-11] (termed SOFI-Airyscan for ease of comparison) and was also more straightforward and required no programming effort. Briefly, multiple Airyscan images were acquired, during which time several cycles Supplementary of light modulation were applied to induce synchronized on- and gradually off-switching of the probe, and finally, SOGO-SOFI processing was performed with the post-deconvolution Airyscan image stacks. We validated the concept by imaging U-2 OS cells expressing Lifeact-Skylan-S fusions. The detailed imaging scheme is illustrated in Supplementary Figure 4a. Specifically, 50 frames of Airyscan images containing 10 cycles of light modulation were recorded. In each cycle, a 405-nm laser was first applied to fully activate Skylan-S, and then, five images were recorded under the illumination of a 488-nm laser. For comparison with SOFI processing, images were acquired by repeating the point-scanning imaging 50 times under continuous irradiation by a 488-nm laser. It is worth noting that a frame number of 50 was chosen according to the preliminary experiment (Supplementary Figure 5), as 50 frames was sufficient for reconstruction with an acceptable quality as compared to a frame number of 500.

Reconstruction results within a small region showed that actin structures resolved by SS-Airyscan exhibited the best continuity and richest details compared to those resolved by confocal microscopy, Airyscan, and SOFI-Airyscan, indicating a high labeling density and a high sensitivity toward weak signals (Figure 4a). The RSP and RSE analysis again confirmed the visual impression that SS-Airyscan SR images had much higher quality and fewer artifacts than SOFI-Airyscan images (Figure 4b–c). The whole cell imaging result further showed that compared to Airyscan, SS-Airyscan notably increased the spatial resolution and suppressed the background (Figure 4d–e) and showed much sharper and thinner structures of actin fibers (Figure 4d–f). Fourier ring correlation (FRC) analysis[32] showed that the spatial resolution of SS-Airyscan was estimated to be 91 nm, which is a 1.5× increase compared to that of Airyscan (140 nm) and is consistent with the theoretical value obtained using the SOFI algorithm.

**Figure 4.**
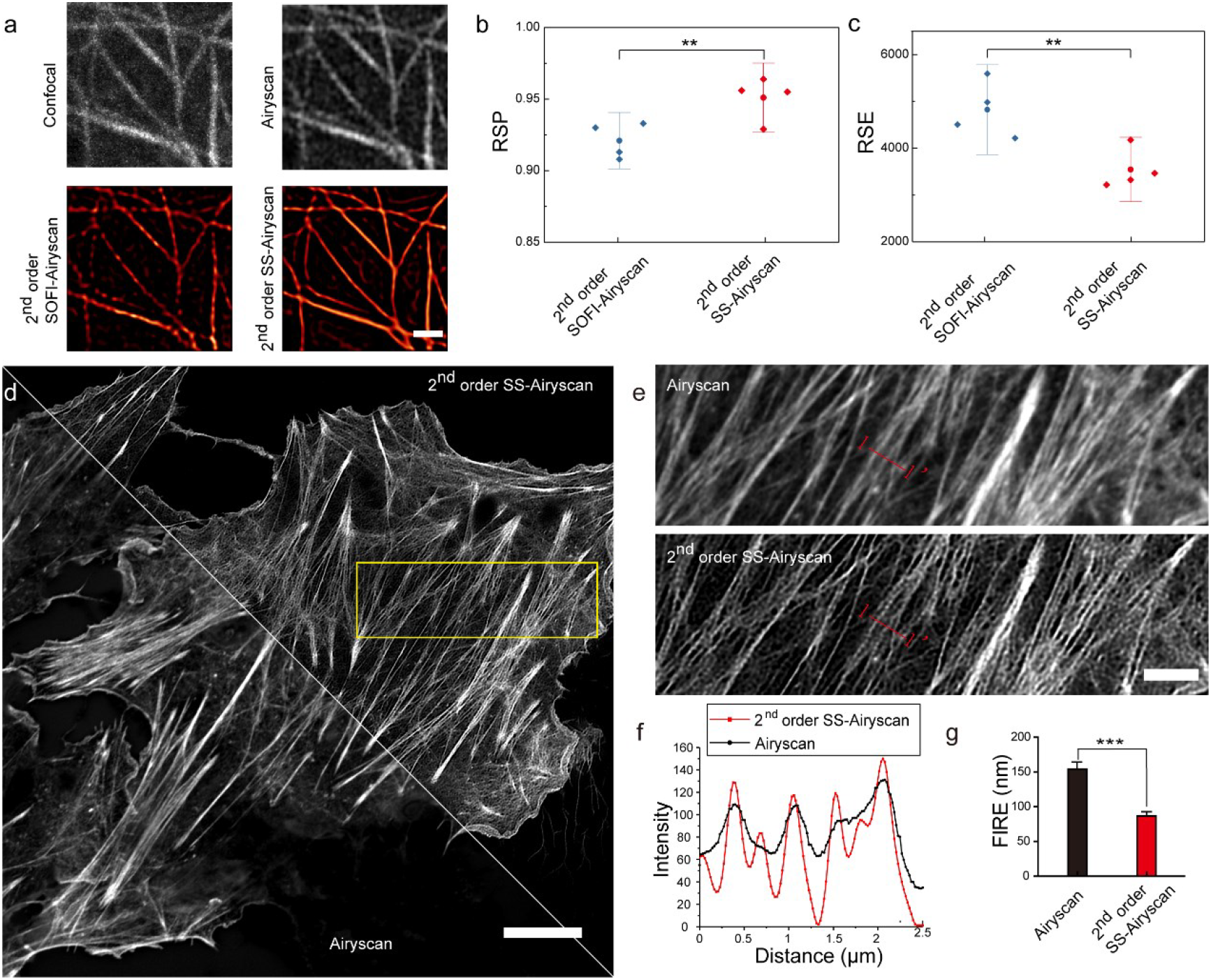
Imaging of F-actin in U-2 OS cells by SS-Airyscan. (a) Images of confocal microscopy, Airyscan, 2^nd^ order SOFI-Airyscan, and 2^nd^ order SS-Airyscan within a small region. (b) Image quality comparison between 2^nd^ order SOFI-Airyscan and 2^nd^ order SS-Airyscan by RSP. (c) Image quality comparison between 2^nd^ order SOFI-Airyscan and 2^nd^ order SS-Airyscan by RSE. (d) Airyscan and 2^nd^ order SS-Airyscan images of a large field-of-view (FOV). (e) Magnified images of the boxed region in (d). (f) Intensity profiles along the cuts 1-1’ in (e). (g) Fourier ring correlation analysis. Scale bars: 1 μm in (a), 10 μm in (d), and 3 μm in (e). Data are reported as mean ± SD. All p values were calculated by the two-tailed Student’s *t*-test, *n* = 5 in (b,c), *n* = 10 in (f), ** indicates p<0.001, *** indicates p<0.0001.

### Resolving SR structures of nucleolar components in cells by dual-color SS-Airyscan

The organization of the nucleolar components (NCs), which mainly contain the fibrillar center (FC), the dense fibrillar component (DFC), and the granular component (GC), is important for the nucleolus to function as a ribosome factory. We first demonstrated the utility of SS-Airyscan for deep imaging by resolving the SR structures of GC and DFC in the nucleolus. The GC marker B23 and the DFC marker Nop56 were double labeled by rsFusionRed3[33] (a red RSFP) and Skylan-S in U-2 OS cells respectively. Dual-color images were acquired with the imaging scheme illustrated in Supplementary Figure 4b. Compared to Airyscan (Figure 5a, 5c), SS-Airyscan (Figure 5b–c) remarkably eliminated the fluorescent background of both channels and more clearly revealed the typical structures of DFC and GC. The signal-to-background ratio (SBR) in both channels was significantly increased by SS-Airyscan compared to Airyscan (Figure 5d–e). This result strongly evidenced the advantage of SS-Airyscan for deep SR imaging.

**Figure 5.**
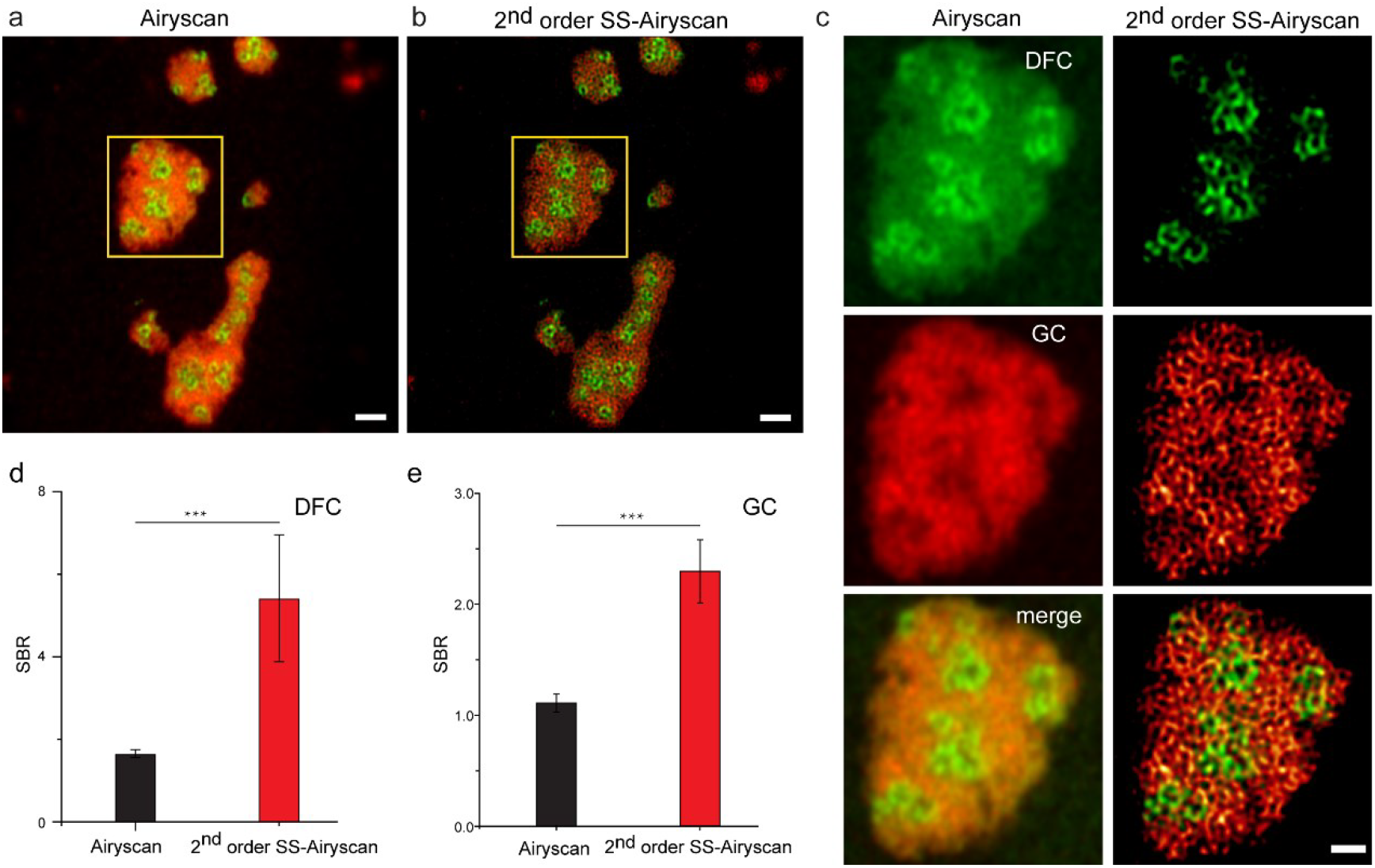
Dual-color imaging of the nucleolar components in cells by SS-Airyscan. (a) Merged dual-color Airyscan images of Nop56 (green) and B23 (red). (b) Merged dual-color 2^nd^ order SS-Airyscan images of Nop56 (green) and B23 (red). (c) Magnified view of the boxed region in (a) and (b). (d) Comparison of signal-to-background ratio (SBR) between the Airyscan and 2^nd^ order SS-Airyscan images in the green channel. (e) Comparison of SBR between the Airyscan and 2^nd^ order SS-Airyscan images in the red channel. Scale bars: 2 μm in (a,b) and 1 μm in (c). Data are reported as mean ± SD. The p values were calculated by the two-tailed Student’s *t*-test, *n* = 14, *** indicates p<0.0001.

### Resolving ER structures in brain slices

Encouraged by the excellent effects of background rejection and resolution enhancement in nucleolar imaging, we attempted to verify the performance of SS-Airyscan in thick tissues. The brain ventricles of newborn mice were infected with calnexin-Skylan-S plasmids by lentivirus. The brains were removed 7 days post infection, embedded in optimal cutting temperature (OCT) medium, and frozen sectioned into slices with a thickness of 50 µm. SS-Airyscan was performed as described before. We found that within a wide range of imaging depths, SS-Airyscan can effectively resolve the porous network structures of ER in the neurons, both in the cell body and in the synapse. Notably, even at a depth of 20.6 µm, the 2^nd^- and 3^rd^-order SS-Airyscan successfully uncovered structural details that were not observed by either Airyscan or SOFI-Airyscan (Figure 6a–b). The spatial resolution of the 2^nd^-order SS-Airyscan measured by FRC analysis was 171 nm, which is significantly higher than that of SOFI-Airyscan (Figure 6c). Notably, the cross-section profiling along the red lines showed the porous structures of ER by SS- Airyscan but not by SOFI-Airyscan (Figure 6d), strongly proving the superiority of SS-Airyscan for deep SR imaging.

**Figure 6.**
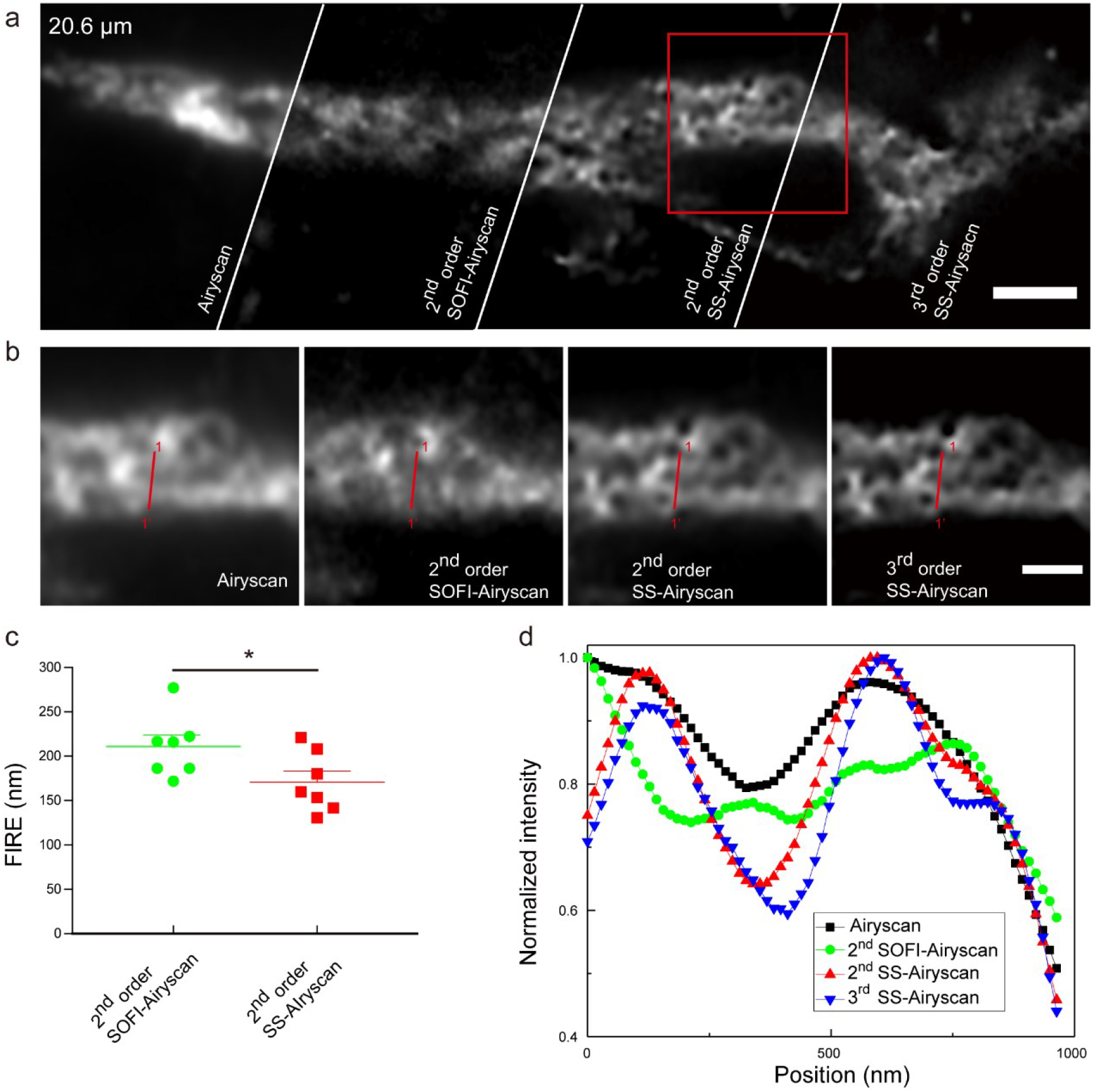
SR imaging of ER in fixed mouse brain slices. (a) Reconstructed images of Airyscan, 2^nd^ order SOFI-Airyscan, 2^nd^ order SS-Airyscan, and 3^rd^ order SS-Airyscan in brain slice at a depth of 20.6 µm. (b) Magnified view of the boxed region in (a). (c) Spatial resolution assessment by Fourier image resolution (FIRE). (d) Intensity cross-section profiles along the red lines indicated by 1 and 1’ in (b). Scale bar: 2 μm in (a) and 1 μm in (b). Data are reported as mean ± SD. All p values were calculated by the two-tailed Student’s *t*-test, *n* = 7, * indicates p<0.05.

## Discussion

In this work, we developed a method, SOGO-SOFI, which is as easy as the conventional pcSOFI but requires 10× fewer raw frames (needing only 20 frames) for reconstruction and gains 10× increased temporal resolution. Moreover, the image quality of SOGI-SOFI, represented here by the high fidelity and few artifacts, is significantly improved compared to pcSOFI. When combined with Airyscan, SS- Airyscan has additive effects on the sectioning capability, background suppression, and spatial resolution enhancement and thus is very attractive for imaging samples that are deep inside cells or have large thickness such as tissue slices. The great capabilities of SS-Airyscan were demonstrated and exemplified by dual-color imaging of nucleolar components in mammalian cells and ER networks in thick brain slices. Meanwhile, the fusion method is still very easy to operate and analyze, reflecting the flexibility of the SOGO-SOFI method.

The superiority of SOGO-SOFI over pcSOFI using only 20 raw frames for reconstruction mainly relies on the light-modulated property of RSFPs. To be specific, on the one hand, light modulation in SOGO- SOFI greatly enhances the probability and frequency of the photoswitching of single RSFP molecules. In contrast, pcSOFI solely relies on the uncontrolled random blinking of the fluorescent probe, thereby wasting the light-modulated activation and deactivation properties of the probes. On the other hand, the modulation scheme of SOGO-SOFI produces a noise-removed signal that can be used for RL deconvolution in the b-SOFI algorithm to improve spatial resolution [34]. The current study exploited the signal changes generated by synchronously activated ON molecules, followed by the unsynchronized disappearance of random fluorescent signals, and performed correlation analysis and pcSOFI. Theoretically, a reverse scheme is also feasible. Such a scheme would involve modulating all the molecules in the off state and then controlling the energy of the 405-nm laser to activate the fluorescent molecules to the on state asynchronously. These asynchronous and randomly generated changes in fluorescence signal could be used for correlation analysis and pcSOFI. This method could also be combined with SOGO-SOFI to further improve the temporal resolution.

SOGO-SOFI should be compatible with a wide range of RSFP probes with different brightness, switching contrast, and kinetics, which has been demonstrated in our dual-color imaging using Skylan- S and rsFusionRed3, two RSFPs with different properties. A probe with high on/off contrast, fast off rate, and high photostability would be beneficial for fast SOGO-SOFI imaging.

SOGO-SOFI is an easy-to-implement method and can be combined with other SR techniques to further increase spatial resolution. In the current study, SOGO-SOFI combined with Airyscan (SS-Airyscan) can increase the image contrast and enhance the resolution of Airyscan by a factor of 1.5 from 140 nm to 91 nm. Due to the slow mechanical conversion between the 405-nm laser for on-switching and the 488-nm laser for off-switching and signal recording of the commercial Airyscan microscope, the temporal resolution of the SS-Airyscan method is currently insufficient for live imaging. It is worth combining SOGO-SOFI with spinning disk confocal-based structured illumination SR techniques in the future to further improve the temporal resolution for live-cell SR imaging. Moreover, SOGO-SOFI could also be combined with various other imaging modalities, such as two-photon, structured illumination, light sheet, and expansion microscopy for further function and application expansions.

## Methods

### Plasmid construction

pSkylan-S-N1 was constructed in a previous study by our lab[13]. To obtain prsFusionRed3-N1, the cDNA sequence of rsFusionRed3 was synthesized (Tsingke, Beijing, China) and inserted into the pEGFP-N1 vector between the AgeI and XhoI restriction sites. To construct pNop56-Skylan-S and pB23-rsFusionRed3, the full-length cDNA sequences of Nop56 and B23 were first cloned from the HeLa cDNA library and then inserted into the pSkylan-S-N1 and prsFusionRed3-N1 vectors, respectively, between the EcoRI and SalI restriction sites. The cDNAs of Lifeact and calnexin were amplified via polymerase chain reaction (PCR) and inserted into pSkylan-S-N1 between the XhoI and BamHI restriction sites. The lentiviral vector expressing ER targeting Skylan-S was constructed by inserting the cDNA of calnexin-Skylan-S into the pCDH CMV vector between the XbaI and NotI restriction sites. All plasmids were sequenced (BGI, China) before further analysis. The restriction enzymes were purchased from New England Biolabs Inc.

### Cell culture, transfection, and fixation

U-2 OS cells were cultured in McCoy’s 5A (modified) Medium (MCMM) (Thermo Fisher Scientific, USA) supplemented with 10% fetal bovine serum (Thermo Fisher Scientific, USA). HEK 293 cells and COS-7 cells were cultured in Dulbecco’s Modified Eagle Medium (DMEM) (Thermo Fisher Scientific, USA) supplemented with glucose (Thermo Fisher Scientific, USA) and 10% fetal bovine serum (Thermo Fisher Scientific, USA). The cells were grown at 37°C with 5% CO_2_. Transient transfections were performed with Lipofectamine™ 2000 (Thermo Fisher Scientific, USA) in 12-well plates following the manufacturer’s protocol when cells reached 80% confluence. At 12 h post transfection, the cells were trypsinized and coated onto slides that were pre-treated with fibronectin and cultured for another 36 h. Fixations were performed with 4% paraformaldehyde and 0.2% glutaraldehyde in PBS buffer (pH 7.4) for 15 min at 37°C, and the products were washed three times with PBS.

### Lentivirus production and transduction

Lentiviral particles were produced by co-transfection of HEK 293T cells with three plasmids—pMD2.G, psPAX2, and pCDH CMV calnexin-Skylan-S—using polyethylenimine (PEI, Polysciences, USA). At 48 h and 72 h post transfection, culture supernatants containing viral particles were collected and clarified with a 0.45-μm membrane filter (Thermo Fisher Scientific, USA) and stored in a deep freezer at −70°C immediately. Titers were determined by limiting dilution. A serial dilution of lentiviral stocks in complete growth medium was prepared in 12-well plates: 1:10^2^, 1:10^3^, 1:10^4^, and 1:10^5^. The plates were incubated at 37°C for 3 d. Then, an inverted fluorescence microscope equipped with a 10× objective and appropriate filter sets was used to visualize the fluorescence so that the total number of colonies per field of view could be counted to calculate the titers. For stereotaxic injection, the lentiviral particles were concentrated by 5× PEG-8000. The supernatant was transferred to a sterile vessel and 1 volume of cold PEG was added to 4 volumes of the supernatant. After 12 h, the supernatant/PEG-8000 mixture was centrifuged at 1500 × g for 30 min at 4°C. Lentiviral pellets were resuspended using cold, sterile PBS at 4°C to 1/100 of its original volume. Then, 1 µL of titrated virus (5 × 10^8^) was injected into the hilus of each BALB/c mouse (1 d). Mouse brains were collected 14 d post injection. Brains were fixed in 4% paraformaldehyde and 0.2% glutaraldehyde in PBS overnight and dehydrated in 30% sucrose for 24 h. After dehydration, the brains were embedded in OCT (Leica, Gernamy) and sectioned on a Cryostat (Leica, CM1900, Gernamy) [35].

### Optical setup and image acquisition

SOGO-SOFI and SOFI were performed on a homemade TIRF microscope composed of an Olympus IX71 body (Olympus, Japan), a 100×, 1.49 NA oil objective (Olympus PLAN APO, Japan), a 1.6× intermediate magnification, and a multiband dichroic filter (Emitter: FF01-446/523/600/647-25 Dichroic: Di01-R405/488/561/635-25×36). We used two lasers (Obis 405-nm LX 50mW, Coherent; Sapphire 488- nm LP 200mW, Coherent, USA) as the light sources, and an acoustic optical tunable filter (AOTF, AA Opto-Electronic, France) to combine, switch, and adjust the illumination power of the lasers. The output lasers were then collimated by an objective lens (L1, CFI Plan Apochromat Lambda 2× N.A. 0.10, Nikon). The light then passed through another lens (L2, AC508-300-A, Thorlabs) and was reflected onto the sample plane. The emitted fluorescence collected by the same objective was passed through a dichroic mirror (DM), an emission filter, and another tube lens (L3) in the microscope body, split by an image splitter (W-VIEW GEMINI, Hamamatsu, Japan), and captured by an sCMOS camera (Prime 95B, Photometrics, USA). The intensity levels of the 405 nm and 488 nm lasers were 18 W/cm^2^ and 39 W/cm^2^, respectively. Airyscan imaging was performed using a commercial confocal microscope (ZEISS, LSM98, Germany) equipped with an Airyscan detector and a Plan-Apochromat 63×/1.4NA oil objective. The pixel time was set to 0.83 μs. The power levels of the 405 nm and 488 nm lasers were set to 47 μW and 106 μW for imaging F-actin, respectively. The power levels of the 405 nm, 488 nm, and 594 nm lasers were set to 31 μW, 159 μW, and 83 μW for imaging nucleoli, respectively. The power levels of the 405 nm and 488 nm lasers were set to 31 μW and 159 μW for imaging brain slices, respectively.

## Statistical analysis

For FRC analysis of SOFI and SOGO-SOFI data, we first split the full image sequence (50 frames) into 10 subgroups as follows: 1–5, 6–10, 11–15, …, 45–50. Then, the odd subgroups were allocated to one subset, and the even subgroups were allocated to another subset. SOFI reconstruction was performed on each subset separately. FRC analysis was carried out on the two subsets by the plugin in Fiji. The image resolution is the inverse of the spatial frequency at which the FRC curve drops below the threshold 1/7.

## Data availability

The datasets generated and/or analyzed in the current study are not publicly available for privacy reasons but are available from the corresponding author on reasonable request.

## Author contributions

P.X. proposed the concept. P.X. and M.Z. designed the experiments. F.X. performed the experiments and analyzed the data. D.P. and H.Y. assisted with sample preparation of the mouse brain slice. W.H. set up TIRF microscopy. P.X. and M.Z. guided the project and wrote the manuscript.

## Supplementary Note

### Data generation for simulation

We generated the data for SOGO-SOFI simulation by defining the number of single molecules at the n^th^ frame, which was composed of the molecules that remained fluorescent from the previous frame and the molecules that were switched on in this frame:

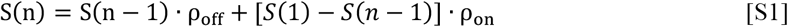

where *S*(1)was set to M.

Assuming that the fluorescent signal of the RSFP ensemble decayed exponentially with the time constant τ_*off*_ within each on-off cycle, and the exposure time is ET, then ρ_off_ can be expressed as

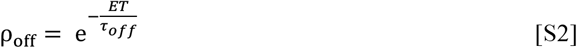

When on-switching and off-switching of the RSFP molecules reach equilibrium, that is

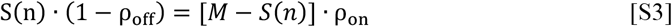

And the on-off contrast of the RSFP probe is given by

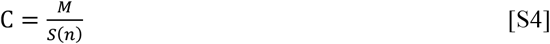

Substituting S3 with S4, we can obtain

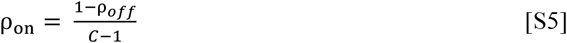

For SOFI simulation, the number of single molecules at the n^th^ frame was defined as

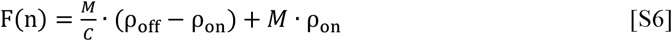

## SI Figures

**Fig. S1.**
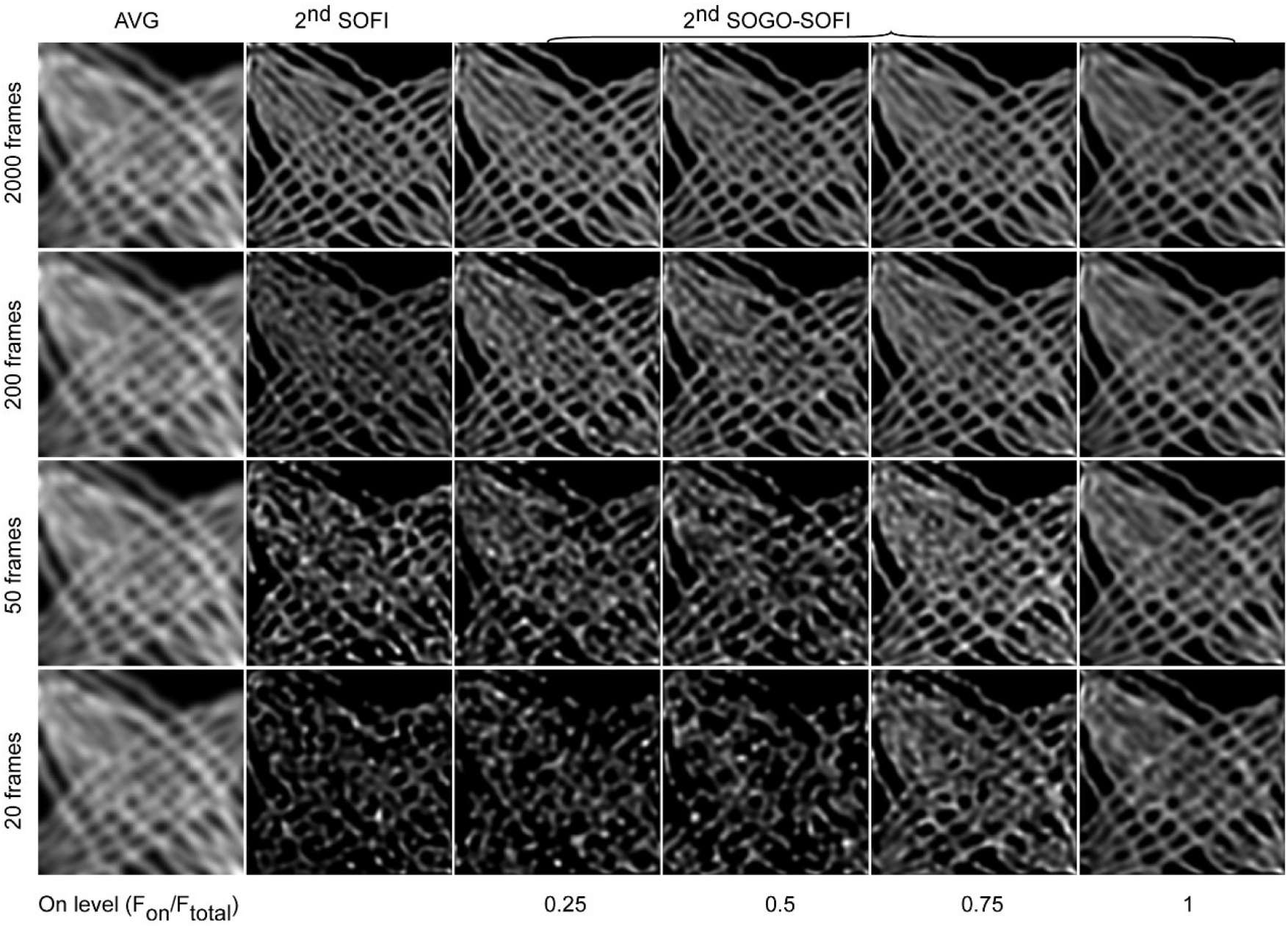
Influence of raw frame numbers and the ratio of ON molecules to all the molecules on the quality of reconstructed SR images by simulation. The numbers of 20, 50, 200 and 2000 raw frames were used for 2^nd^ SOFI and 2^nd^ SOGO-SOFI reconstructions. The ratios of 0.25, 0.5, 0.75 and 1 were used in 2^nd^ SOGO-SOFI for raw image generation. AVG: averaged image from different raw frames.

**Fig. S2.**
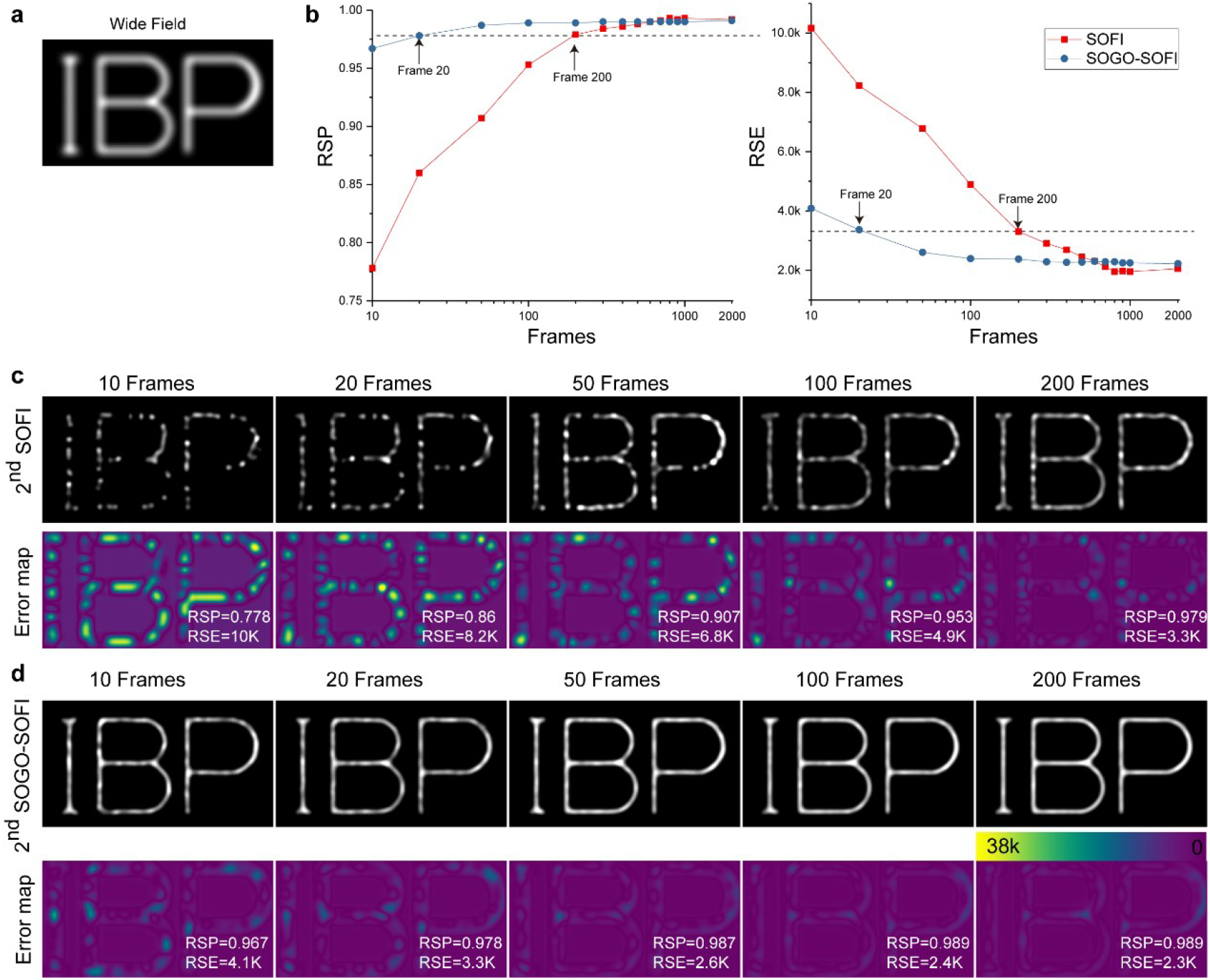
Determination of the minimal frame numbers for SOGO-SOFI reconstruction. (a) The simulated wide field image. (b) Comparison of RSE and RSP between SOFI and SOGO-SOFI with different frame numbers in log transformed space. (c) Reconstructed SR images and the corresponding error maps of SOFI with different raw frame numbers. (d) Reconstructed SR image and the corresponding error map of SOGO-SOFI with different raw frame numbers.

**Fig. S3.**
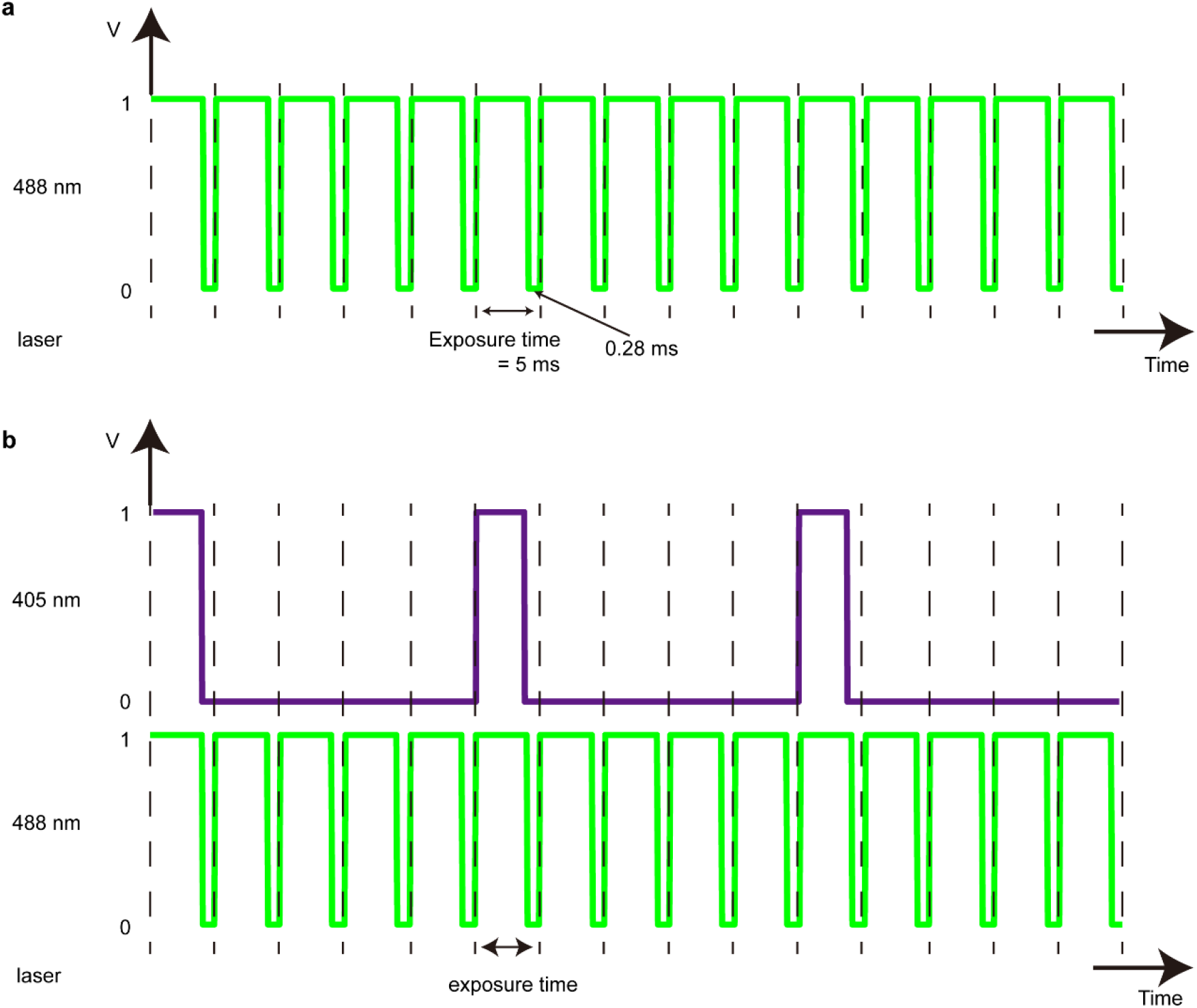
Imaging acquisition schemes of SOFI (a) and SOGO-SOFI (b).

**Fig. S4.**
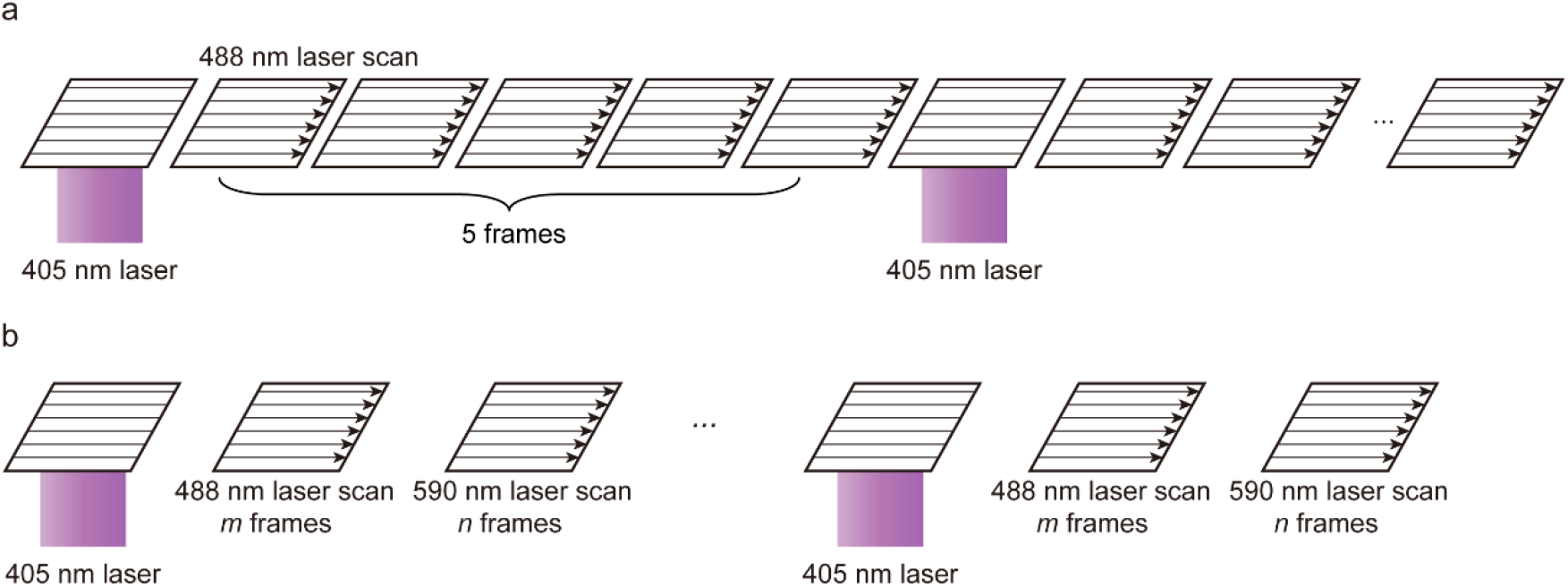
Imaging acquisition schemes of single-color (a) and dual-color (b) SS-Airyscan.

**Fig. S5.**
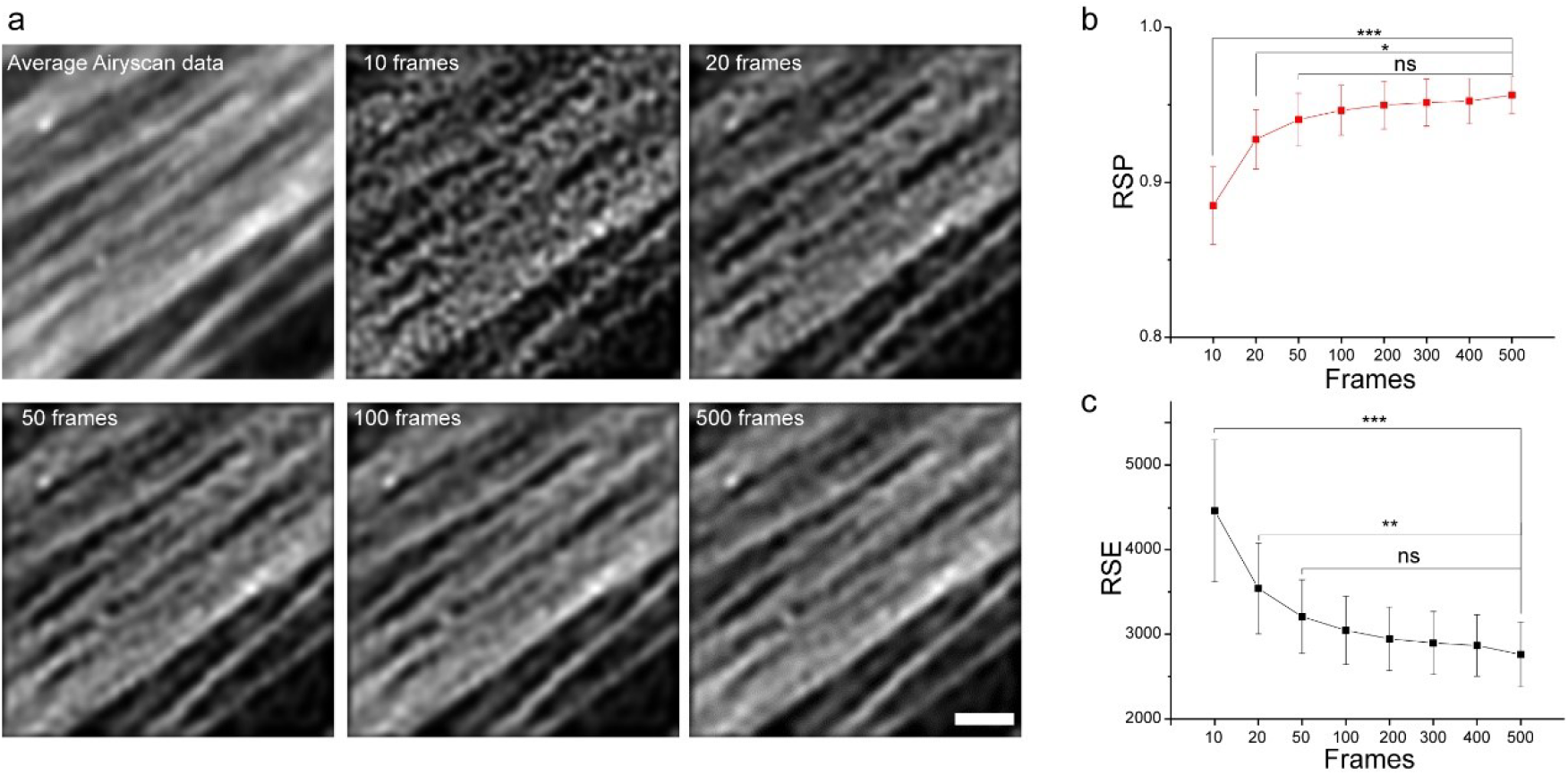
Determination of the minimal frame number for SS-Airyscan reconstruction. (a) Averaged Airyscan image and reconstructed SR images of SS-Airyscan with different frame numbers. (b, c) Comparison of RSP and RSE depending on the frame numbers.

